# Comparison of computational methods for imputing single-cell RNA-sequencing data

**DOI:** 10.1101/241190

**Authors:** Lihua Zhang, Shihua Zhang

## Abstract

Single-cell RNA-sequencing (scRNA-seq) is a recent breakthrough technology, which paves the way for measuring RNA levels at single cell resolution to study precise biological functions. One of the main challenges when analyzing scRNA-seq data is the presence of zeros or dropout events, which may mislead downstream analyses. To compensate the dropout effect, several methods have been developed to impute gene expression since the first Bayesian-based method being proposed in 2016. However, these methods have shown very diverse characteristics in terms of model hypothesis and imputation performance. Thus, large-scale comparison and evaluation of these methods is urgently needed now. To this end, we compared eight imputation methods, evaluated their power in recovering original real data, and performed broad analyses to explore their effects on clustering cell types, detecting differentially expressed genes, and reconstructing lineage trajectories in the context of both simulated and real data. Simulated datasets and case studies highlight that there are no one method performs the best in all the situations. Some defects of these methods such as scalability, robustness and unavailability in some situations need to be addressed in future studies.

## 1 Introduction

High-throughput RNA sequencing technology has been successfully applied to quantify transcriptome profiling. However, it usually takes advantages of millions of cells to quantify gene expression, which is insufficient for studying heterogeneous systems, e.g. embryo development, brain tissue formation and tumor differentiation. Single-cell RNA-sequencing (scRNA-seq) technology was first reported by Tang in 2009 [1], and gained widespread attentions until 2014 when the protocols become easily accessible. Currently, many efficient sequencing technologies are constantly emerging, such as Smart-seq, Dropseq, CEL-seq, SCRB-seq and the commercial device 10X chromium3.

scRNA-seq has revealed distinct heterogeneous of individual cells within a seemingly homogeneous cell population or tissue, and provided insights into cell identity, fate and function [2], [3]. Many computational methods from traditional bulk RNA sequencing (bulk-RNAseq) data may be useful for analyzing the scRNA-seq data. However, there are some differences between them. One main difference from bulk-RNAseq is that scRNA-seq takes each cell as a sequencing library. However, the amount of mRNAs in one cell is tiny (about 0.01–0.25pg), and it has up to one million fold amplification. A low starting amount makes some mRNAs are totally missed during the reverse transcription and cDNA amplification step, and consequently cannot be detected in the latter sequencing step. This phenomenon is the so-called ‘dropout’ event, which suggests that a gene is observed in one cell with moderate or high expression level, but not detected in another cell [4], [5].

There are also missing values in bulk-RNAseq or microarray data. Many imputation methods have been proposed to address this issue [6], [7], [8]. For example, Kim *et al*. proposed a local least squares imputation method named LLSimpute [6], which imputes each missing value with a linear combination of similar genes. However, these imputation methods may be not directly applicable to scRNA-seq data. As bulk-RNAseq measures the average gene expression, while scRNA-seq can detect gene expression at single cell resolution. There would be more data fluctuation in scRNA-seq than that in bulk-RNAseq. Moreover, scRNA-seq data is much sparse than bulk-RNAseq data.

Considering the famous Netflix problem in the area of recommendation system: as users only rate a few items, one would like to infer their preference for unrated ones. Obviously, only a few factors affect an individual’s preference. Thus, the user-rating data matrix should be in low-rank. Interestingly, the single cell gene expression data matrix should also be in low-rank as the limited cell subpopulations and distinct homogeneity in a cell population. Thus, the low-rank matrix completion method (Low-rank) [9], [10] can also be applied to the scRNA-seq data imputation problem.

Several imputation methods designed specifically for scRNA-seq data have been proposed in recent studies. BISCUIT adopts a Dirchlet process mixture model to iteratively normalize, impute data, and cluster cells by simultaneously inferring parameters of clustering, capturing technical variations (e.g. library size), and learning cluster-specific co-expression structures. Therefore, BISCUIT gives out a normalized and imputed data matrix. However, BISCUIT is a MCMC-based method, which costs lots of time to implement [11]. scUnif is a unified statistical framework for both single cell and bulk RNA-seq data [12]. However, scUnif is a supervised learning method and it needs predefined cell type labels that are often unknown. MAGIC is a Markov affinity-based graph imputation method, which weights other cells by a Markov transition matrix [13]. However, it also imputes counts that are not affected by dropout. Therefore, it may introduce new bias into the data and possibly eliminate meaningful biological variations. scImpute separates genes into two gene sets for cell *j* (unreliable and reliable categories: *A_j_*, *B_j_*) based on a dropout probability, which is estimated by a mixture model [14]. scImpute imputes *A_j_* by treating *B_j_* as gold-standard data. And a weighted LASSO model is used on genes in *B_j_* across other cells to find similar cells. Then *A_j_* is imputed by a linear regression model with the most similar cells. scImpute could distinguish the dropout zeros and real zeros. However, scImpute assumes that each gene has an overall dropout rate, while it has been verified that the dropout rate of a gene is dependent on many factors such as cell types, RNA-seq protocols [4]. LASSO tends to select just one cell if there are many cells highly correlated with cell *j* [15], which may ignore some useful information. DrImpute is an ensemble method, which is designed based on a consensus clustering method [16] for scRNAseq data. In other words, it performs clustering for many times and conducts imputation by the average value of similar cells [17]. SAVER is a global method [18], which recovers the true expression for each gene in each cell by a weighted average of the observed count and the predicted value. The predicted value is estimated by the observed expression of some informative genes in the same cell. We summarize these methods in Table 1. They can be classified into two categories according to wether it is a Bayesian-based method. BISCUIT, scUnif and SAVER are three Bayesian-based ones. They can also be categorized into local or global methods according to how the imputation information is used from the observed data. MAGIC, Low-rank, BISCUIT, scUnif and SAVER are global methods, while the remaining ones including LLSimpute, scImpute and DrImpute are local ones.

**TABLE 1.**
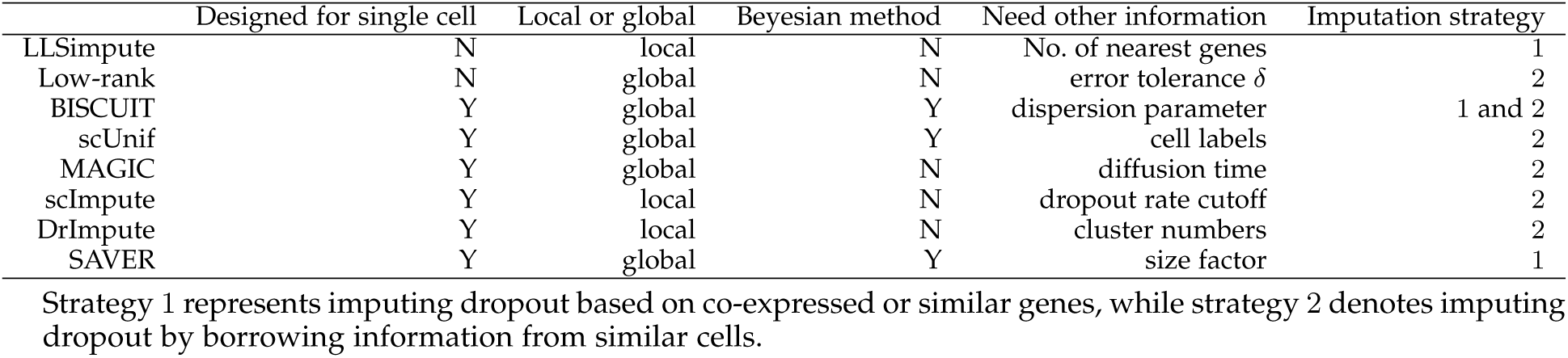
Summary of the eight imputation methods

In this paper, we comprehensively compared and evaluated these imputation methods with both simulated and real data. The rest of this paper is organized as follows. In section 2, we describe more details about the eight imputation methods, datasets used in this study, and the evaluation strategies employed to make comparison. In section 3, we present the performance of these imputation methods extensively. In section 4, we summarize this study and discuss potential directions for imputing scRNA-seq data in future.

## 2 Methods and Materials

### 2.1 Method details

In the followings, the eight imputation methods are described in detail. The observed gene expression data of *m* genes across *n* cells is denoted as *G* ∈ *R^m^*^×^*^n^*, which is obtained from the normalized gene count data (see 2.5). LLSimpute is designed based on a linear regression model, which divides cells into two groups (*C_i_* and *D_i_*) for gene *i*. *C_i_* stores cells in need of imputation (i.e., *C_i_* = {*a*|*G*(*i*, *a*) = 0}), while cells in *D_i_* have reliable gene expression. Suppose there are *q* missing values for gene *i*, it finds the *K*-nearest neighbor gene vectors for gene *i* based on values in *D_i_*, which is represented as 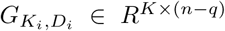. Let 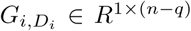 denote gene expression of gene *i* across cells in *D_i_*, and 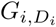 is represented as a linear combination of rows of 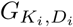 by:

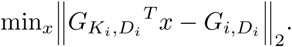

Then the missing values of gene *i* denoted by 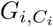 are imputed by 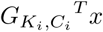, where 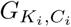 represents gene expression values of genes *K_i_* across cells *C_i_*.

Low-rank method adopted here [9] supposes that gene expression matrix *X* without dropout events is low-rank and can be approximated by its nuclear-norm, which is its convex envelope. The model is summarized as follows,

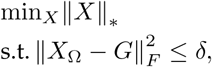
where *X* is the imputed gene expression matrix, *G* is the observed one, Ω is the observed space, and *δ* is error tolerance between the imputed data and the observed one.

BISCUIT is the first approach specially designed for scRNA-seq imputation. Let *X* ∈ *R^m^*^×^*^n^* denote the log-transformed count matrix with pseudo count 1. BISCUIT assumes that each gene expression vector x*_j_* of each cell *j* follows a Gaussian distribution and the likelihood of *x_j_* is *x_j_* ~ *N*(*α_j_μ_k_, β_j_*Σ*_k_*), where *α_j_*, *β_j_* are cell-dependent scaling factors, *μ_k_*, Σ*_k_* are the mean and covariance of the kth mixture component, respectively. The conjugate prior of each *μ_k_* is normal, Σ*_k_* is Wishart, *α_j_* is normal, and *β_j_* is Inverse-gamma. The ideal gene expression of cell *j* after removing technical variations is denoted as 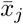, 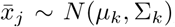, which is the *j*th column of the recovered matrix 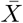 by BISCUIT.

scUnif is a unified framework and it incorporates single cell and bulk data together to obtain more accurate expected relative expression level *E* ∈ *R^m^*^×^*^k^*, where *m* represents the number of genes, *k* denotes the number of cell types, and the sum of each column of *E* is 1. We merely depict the model on scRNA-seq data due to the lack of corresponding bulk-RNAseq data. The gene expression vector for each cell *j* denoted as *G._j_* is assumed to follow a multinomial distribution with probability vector *p_j_* and the number trials *R_j_*, which approximates sequencing depth 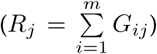. The *i*th entry of *p_j_* is computed as follows,

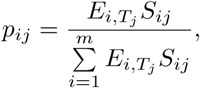
where *S* is a binary variable which represents dropout status and follows a Bernoulli distribution with the observed probability *π_ij_* of gene *i* in a single cell *j*. *π_ij_* is modeled as a logistic function of expected relative expression 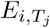 as follows, 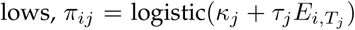, where *T_j_* ∈ {1, 2, …, *k*} is the cell type of cell *j*, *κ_j_* and *τ_j_* are parameters following 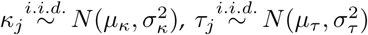 for *j* = 1, 2,…,*n* and *μ_κ_*, *σ_κ_*, *μ_τ_*, *σ_τ_* are parameters. Finally, the expected relative expression profile *E* is inferred by scUnif. Therefore, the dropout value of gene *i* in a single cell *j* is imputed by the multiplication of 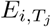 and sequencing depth of cell *j*.

SAVER models the observed count value of gene *i* in a single cell *j* by 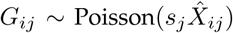, where *s_j_* is a cell-specific size factor and 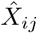 is the normalized true expression level of gene *i* in cell 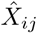 is recovered with the help of *µ_ij_* with a dispersion parameter *φ_i_*, where *µ_ij_* is predicted from the expression of other genes in the same cell. To account for the recovery uncertainty, a Gamma prior is placed on 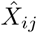: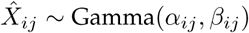, where *α_ij_* and *β_ij_* are the reparam-eterization of *µ_ij_* and *φ_i_*. Then the posterior distribution of 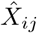 is also gamma distributed and the posterior mean is:

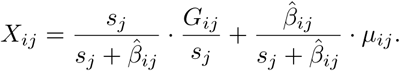

MAGIC leverages the shared information of similar cells to impute missing values. Firstly, MAGIC computes cell-cell distance matrix denoted as *D_dist_* based on Euclidian distance. Then it converts *D_dist_* to an affinity matrix *F* using an adaptive Gaussian kernel. After that, MAGIC transforms *F* to a Markov transition matrix *M* by symmetrizing and normalizing each row of *F*. Finally, MAGIC obtains imputed data matrix *X* by information flows from similar cells in terms of 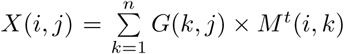, where *G* is the observed gene expression matrix, and *t* represents the diffusion time. Smaller *t* could not capture effective gene structure information, while larger *t* will result in over-smoothing and loss of information after imputation. Therefore, choosing an optimal diffusion time *t* is a key component of MAGIC.

scImpute models the expression levels of gene *i* as the following mixture model 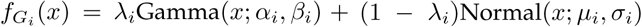, where *λ_i_* represents gene’s dropout rate, *α_i_*, *β_i_* are parameters of Gamma distribution, and *µ_i_*, *σ_i_* are parameters of normal distribution. The dropout probability is computed as follows,

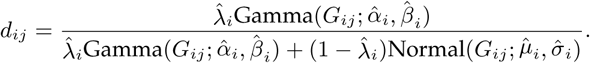

Then scImpute divides genes for each cell *j* into *A_j_*: {*i*: *d_ij_* ≥ *t*} and *B_j_*: {*i*: *d_ij_ <t*}, and genes in *A_j_* are treated as dropout genes in cell *j*, while genes in *B_j_* are thought to have accurate values. scImpute imputes dropout values cell by cell. It constructs a weighted lasso regression model on gene expression of *B_j_* to select similar cells. Then gene expression of *A_j_* is imputed by the ordinary least square linear regression model on similar cells. The threshold value *t* is a key parameter of scImpute.

DrImpute computes the distance of cells using Spearman and Pearson correlations. Then it performs *K*-means clustering on the first 5% principal components of similarity matrix converted from the distance matrix with varied cluster number *k*. Therefore, clustering results *C*_1_*,C*_2_*,…,C*_2_*_k_* are obtained, where the first *k* clusters are based on Spearman correlation distance and the last *k* clusters are based on Pearson correlation distance. The expected value of the dropout is computed by *E*(*x_ij_* |*C_l_*)= mean(*x_ij_* |*x_ij_* are in the same cell group in clustering *C_l_*). Finally, DrImpute imputes the dropout value by averaging the multiple expected values across all clustering results.

### 2.2 Evaluation strategies

We evaluate the performance of imputation methods from two angles. Firstly, the imputed value should be similar to the original value, which can be evaluated in the formation of mean squared error (MSE) and Pearson correlation coefficient (PCC). Secondly, a good recovery method should preserve the biological structures of the data (e.g. cell-type clusters, differently expressed genes (DEGs), and cell differentiation directions). Methods of dimension reduction, clustering, detecting DEGs, and reconstructing pseudotime trajectory for analyzing scRNA-seq data have been developed [4], [16], [19], [20], [21], [22], [23], [24]. Some of these methods consider or impute dropout events, while others do not. In this study, we compared methods considering dropout events or not to study the impact of imputation methods on scRNA-seq analyses.

High level of noise in both technical and biological aspects with large gene or cell dimensions makes scRNA-seq data analyses difficult. Thus, dimension reduction is essential for data visualization and analysis. PCA [25] and tSNE [26] are two commonly used dimension reduction methods. Recently, Zero Inflated Factor Analysis (ZIFA) has been developed to reduce dimensions of scRNA-seq data, which considers dropout events by modeling dropout rate [19]. CIDR is a dimension reduction and clustering method, which incorporates imputation procedure meanwhile [20]. However, the imputation value of a gene in a cell is dependent on another cell it pairs up. In this study, we visualized data by PCA, tSNE, ZIFA and CIDR.

*De novo* discovery of cell-type clusters is one of the most promising application of scRNA-seq. SC3 is a consensus clustering method with a series of ranks based on spectral clustering for analyzing scRNA-seq data [16]. This method does not address the dropout events. A multi-kernel learning based method named SIMLR has been suggested to be robust to dropout events, which also doesn’t consider to addressing dropout events [21]. We implement SC3, SIMLR and *k*-means with the first two tSNE dimensions (tSNE+kmeans) on raw data, original data (if available), and imputed data, respectively.

The negative binomial model fits bulk-RNAseq data very well and several statistical methods have been designed based on this model. For example, edgeR is one of such methods designed for differential expression analysis [27]. However, a raw negative binomial model does not fit single cell read count data well due to dropout. Zero-inflated negative binomial models have been proposed (e.g. SCDE, MAST) for detecting DEGs from scRNA-seq data [4], [22]. SCDE models gene-specific expression with the mixture of a poisson and negative binomial model, and provides the posterior probability of being DEG for each gene between two biological conditions [4]. MAST uses a Gaussian generalized linear model describes expression condition on non-zero expression and tests differential expression rate between groups [22]. We detected DEGs by edgeR on raw data, original data (if available) and imputed data, and MAST, SCDE on raw data respectively.

scRNA-seq has already been used to study cellular transitions between different states. Monocle 1 and Monocle 2 are two widely used methods to deduce the underlying developmental trajectories [23], [24]. However, it does not address dropout. In this study, we applied Monocle 1 and Monocle 2 on raw data and imputed data respectively.

### 2.3 Simulated datasets

Splatter and PowsimR are two R Bioconductor packages proposed recently for reproducible and accurate simulation of scRNA-seq data [28], [29]. PowsimR is designed to simulate and evaluate differential expression for bulk and single cell RNA-seq data. Here we adopted Splatter to generate five scRNA-seq datasets including single or multiple cell populations, cells along a differentiation path, and cells in various batches with predefined or estimated parameters (Table 2). Firstly, we simulated an observed count matrix with 1000 genes and 100 cells in a single population (dataset 1), and set the *dropout.shape* parameter ranging from 0.05 to 0.25 in step of 0.05 resulting in data with increasing dropout ratios. Then we simulated two datasets with multiple subpopulations. One dataset (dataset 2) was of small size with 150, 50, 50 cells in each group and the left parameters were as follows, nGenes = 500, mean.shape = 0.3, mean.rate = 0.02, de.prob = *c*(0.05, 0.02, 0.03), de.facLoc = 0.1, and de.facScale = 0.4. Another one (dataset 3) was of large size and the parameters were: nGenes = 1000, groupCells = *c*(240, 120, 100, 20, 370, 150), de.facLoc = 0.1, and de.facScale = 0.4. Moreover, we simulated a dataset (dataset 4) with 1000 genes and 100 cells. The cells were generated along a differentiation path with default parameters. Finally, we simulated a dataset (dataset 5) with 2000 genes and 100 cells with two groups in two batches with group.prob = c(0.5, 0.5) and other default parameters.

**TABLE 2.** Summary of both simulated and real datasets used for systematic evaluation

|  | Simulated data |  |  |  |  | Real data |  |
| --- | --- | --- | --- | --- | --- | --- | --- |
| Dataset | 1 | 2 | 3 | 4 | 5 | mECS | Mouse IFE |
| Data size | 1000*100 | 500*200 | 1000*1000 | 1000*100 | 2000*100 | 8989*182 | 26165*1529 |
| Cell clusters | 1 | 3 | 6 | a path | 2 (batch effects) | 3 | a path (3 stages) |

### 2.4 Real datasets

We adopted two real biological single cell datasets for this evaluation study (Table 2). The mESC dataset was obtained from a controlled study that explored the effect of cell cycle on gene expression level of individual mouse embryonic stem cells (mESCs) [30]. This data has been used for visualizing, reducing dimensions and clustering single cells in a previous study [21]. We obtained the preprocessed data by this study and there were 182 mouse embryonic stem cells (mESCs) in three cell cycle stages (G1, S and G2M) marked by fluorescence-activated cell sorting [30].

The mouse IFE data was obtained from Gene Expression Omnibus with GSE67602. This data consists of 25932 genes and 536 cells, which were used to reconstruct interfollicular epidermis (IFE) cell differentiation in a previous study [31]. We removed genes that were expressed in less than 5 cells, and kept 13689 genes in the final dataset.

### 2.5 Data preprocessing

For all datasets except the mouse IFE data, if a gene was expressed in less than two cells, it was removed. We normalized the count values by a global normalization method with being divided by library size and multiplied by mean library size across cells. Then the normalized values were log-transformed. scImpute, BISCUIT and scUnif can process this transformation automatically.

## 3 Results

### 3.1 Recover gene expression of a homogenous cell population

We applied the eight methods to the simulated dataset 1, which is a homogenous data with varying ratios of dropout events. We can clearly see that the MSE values increase and PCC values decrease with the ratio of dropout events increasing. Low-rank shows the best performance than other methods with the smallest MSE and the largest PCC values. scUnif and LLSimpute also have better performance, while scImpute has large fluctuations when the ratio of dropout events increases (Figure 1A and 1B). We compared the imputed data with the original one in zero and non-zero spaces respectively on the data with *dropout.shape* = 0.05 (Figure 1C). In the zero space, LLSimpute and Low-rank recover values similar to the original ones. While scImpute imputes the missing values with distinct dispersion. There is a clear linear relationship between the imputed values of scUnif and the original ones. And the imputed values of scUnif is smaller than the corresponding original ones. That is because the imputed value equaled the multiplication of the relative profile and the sequencing depth of the corresponding cell, which is usually unknown and estimated by the sum of raw counts of the corresponding cell. As the existence of dropout, the estimated sequencing depth is lower than the real one. The imputed value of dropout are near zeros by the remaining methods on this data. Among these methods, Low-rank, MAGIC, BISCUIT and SAVER could change the observed values in principle. scImpute may change some observed values, whose dropout probability are smaller than a certain threshold. In the observed non-zero space of the homogenous data, Low-rank, MAGIC and SAVER can recover the original values well, while BISCUIT recovers them with some fluctuations.

**Fig. 1.**
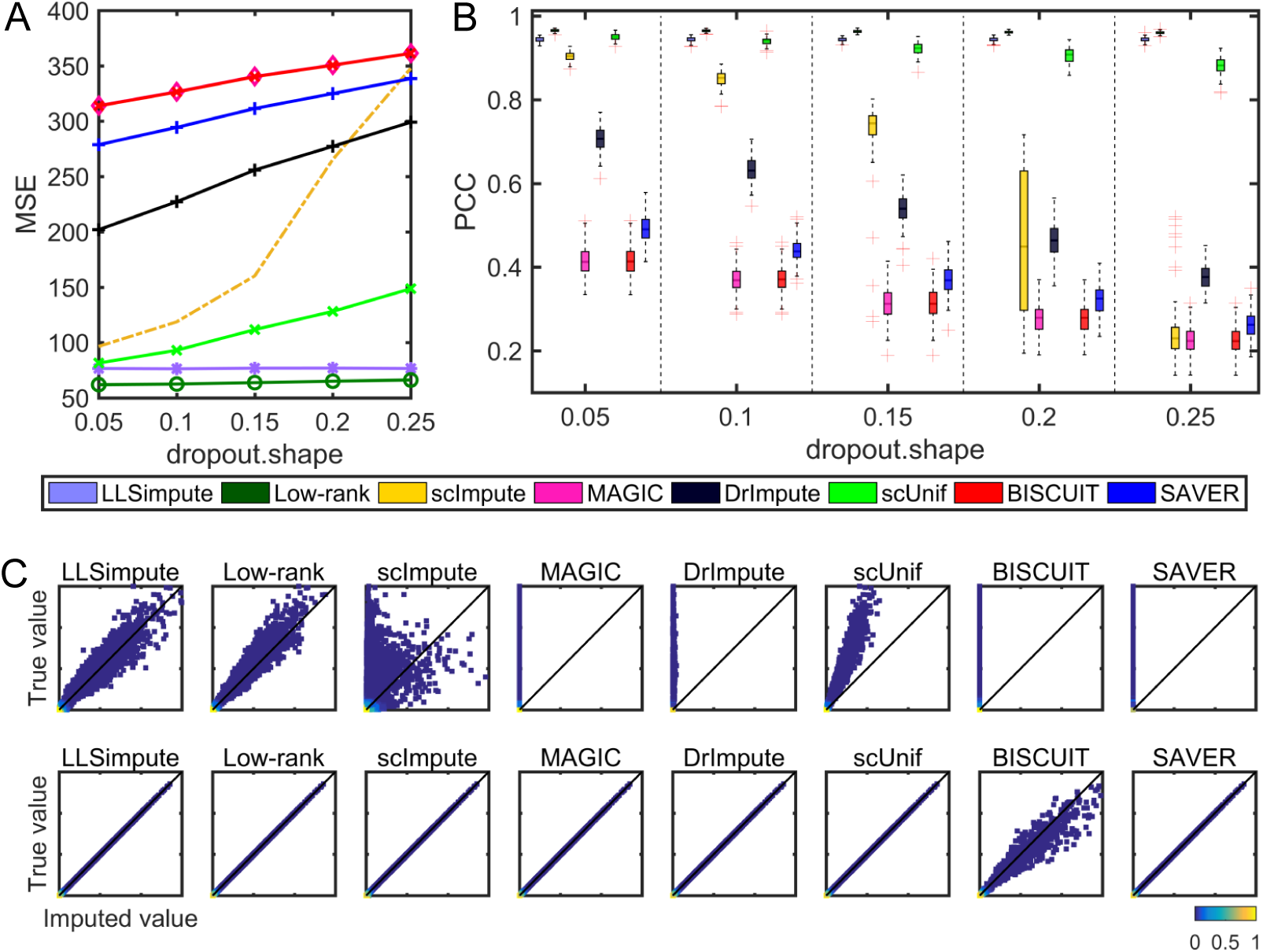
Direct evaluation of the eight imputation methods on simulated dataset 1. (A) MSE values varies across various dropout ratios. (B) PCC values of all single cell pair computed between the imputed data and the original one. (C) Density plot of the imputed values versus the original ones in the zero space (top) and the observed non-zero space (bottom), respectively.

### 3.2 Recover gene expression of the heterogenous scRNA-seq data

We simulated two heterogenous scRNA-seq data with small size (dataset 2) and large size (dataset 3) respectively. LLSimpute shows the largest MSE and the smallest PCC values among all methods on datasets 2 and 3. LLSimpute imputes many true near zero values with large ones. These phenomenon demonstrates that LLSimpute is not applicable to scRNA-seq data directly due to the existence of heterogeneity and sparsity. DrImpute, scImpute, scUnif (given label information), SAVER and Low-rank have better performance in terms of MSE and PCC values (Figure 2A and 2B; Figure 3A and 3B). The Bayesian-based methods took more time to conduct computation (Figure 2C and Figure 3C). Since the negative values imputed by LLSimpute, Low-rank, MAGIC and BISCUIT are meaningless, we set these values to be zeros. BISCUIT and SAVER impute the dropout events by near zero values. In the observed non-zero space, MAGIC and BISCUIT estimate the observed values with relatively large fluctuations, while other methods recover them well.

**Fig. 2.**
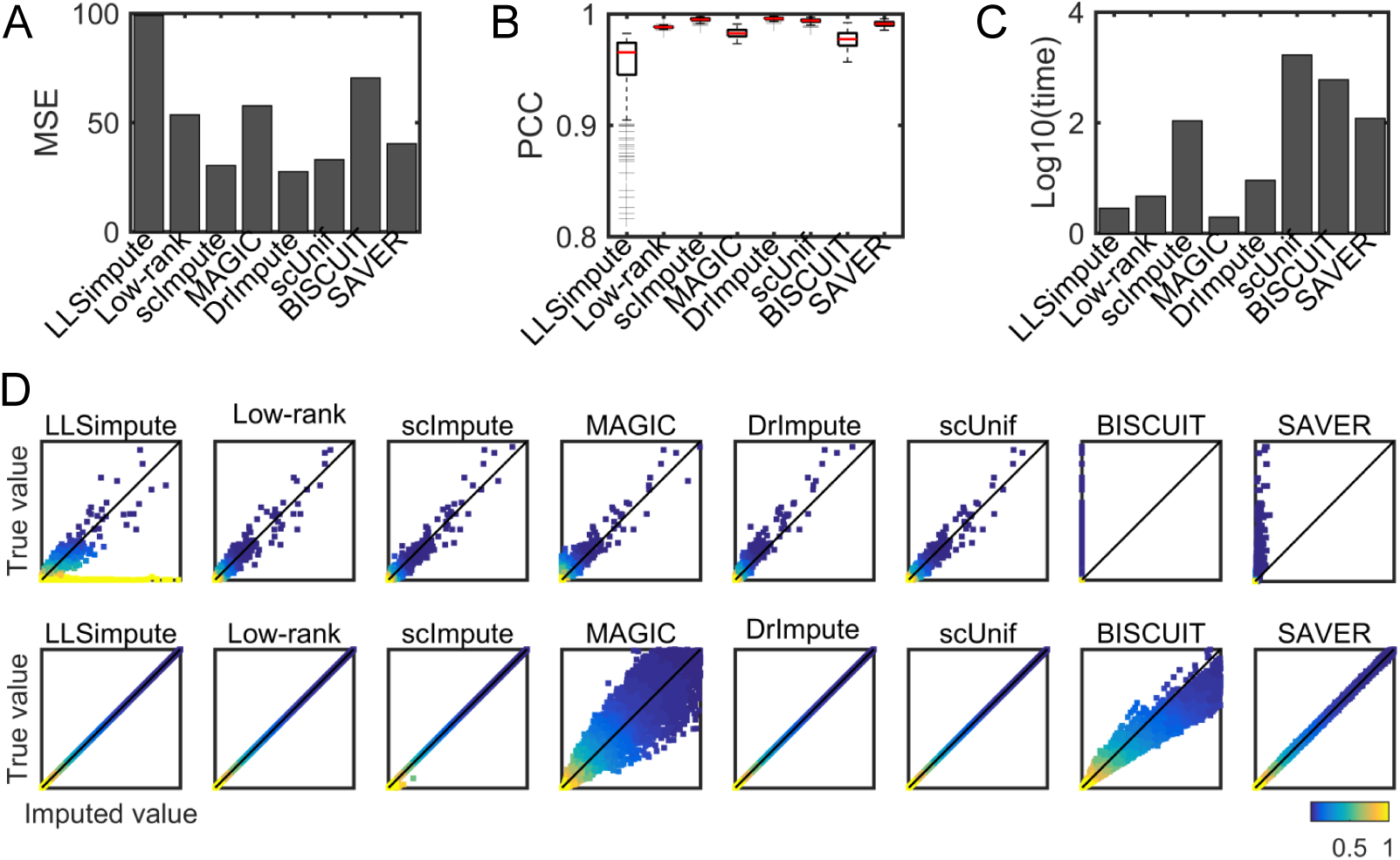
Direct evaluation of the eight imputation methods on simulated dataset 2. (A) MSE values of each imputation method. (B) PCC values of all single cell pair computed between the imputed data and the original one. (C) Computational time (seconds) of running each imputation method. (D) Density plot of the imputed values versus the original ones in the zero space (top) and the observed non-zero space (bottom), respectively.

**Fig. 3.**
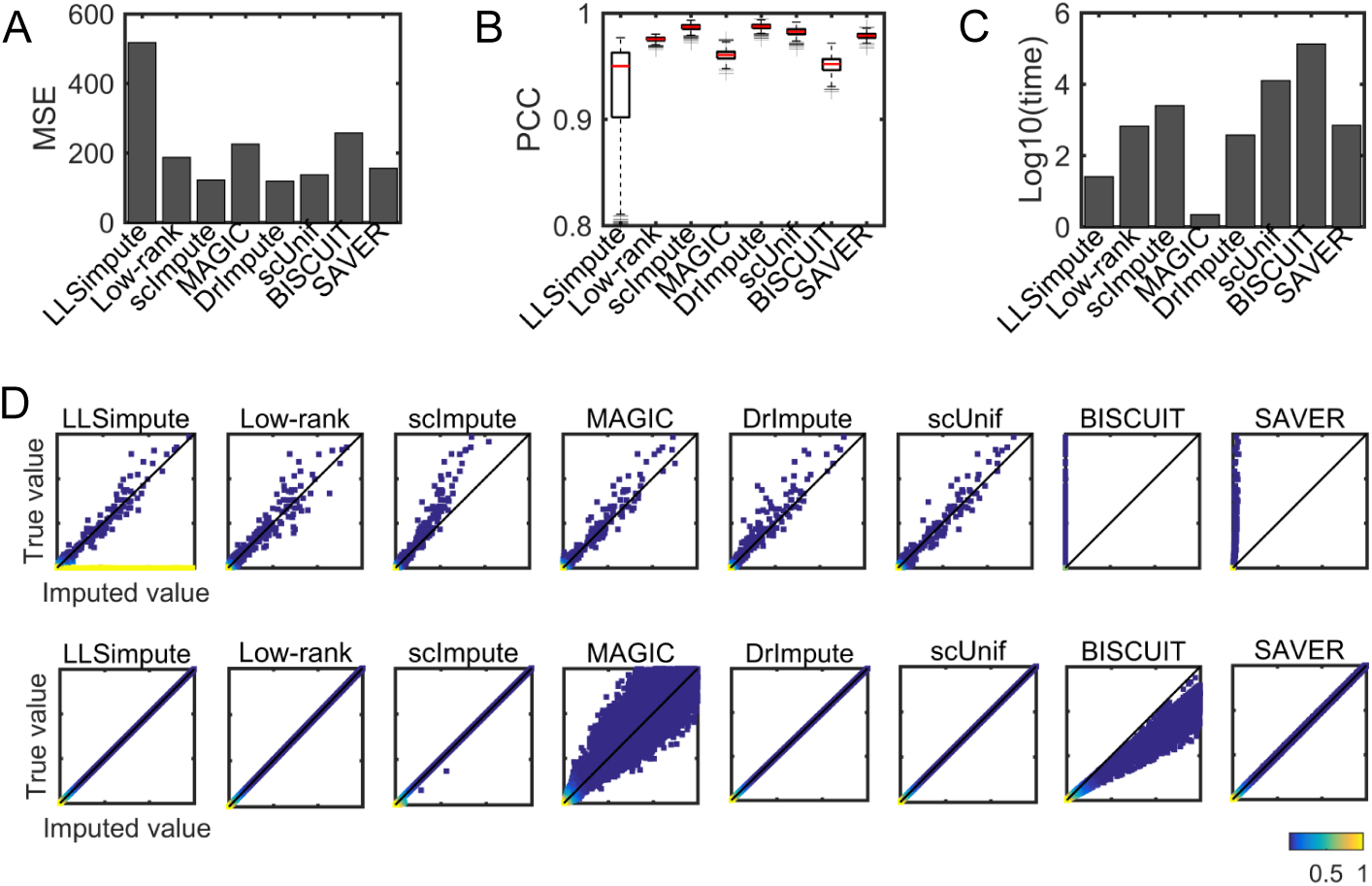
Direct evaluation of the eight imputation methods on simulated dataset 3. (A) MSE values of each imputation method. (B) PCC values of all single cell pair computed between each cell of the imputed data and the original one. (C) Computational time (seconds) of running each imputation method. (D) Density plot of the imputed values versus the original ones in the zero space (top) and the observed non-zero space (bottom), respectively.

We also simulated a heterogenous scRNA-seq data (dataset 5) with batch effect. By PCA visualizing, we can see that the cells in this raw data are mixed together, while the cells in the full data are separated due to batch effects. Interestingly, the batch effects are stronger than group effects with the former represented as the first component, while another as the second component (Figure 4B). Low-rank performs the best on this data with the largest PCC and smallest MSE values. Both imputation methods recover non-zero values well (Figure 4A). This might because that many similar cells were clustered together as illustrated by PCA visualization on the normalized data. However, BISCUIT and SAVER might fail to capture batch effect.

**Fig. 4.**
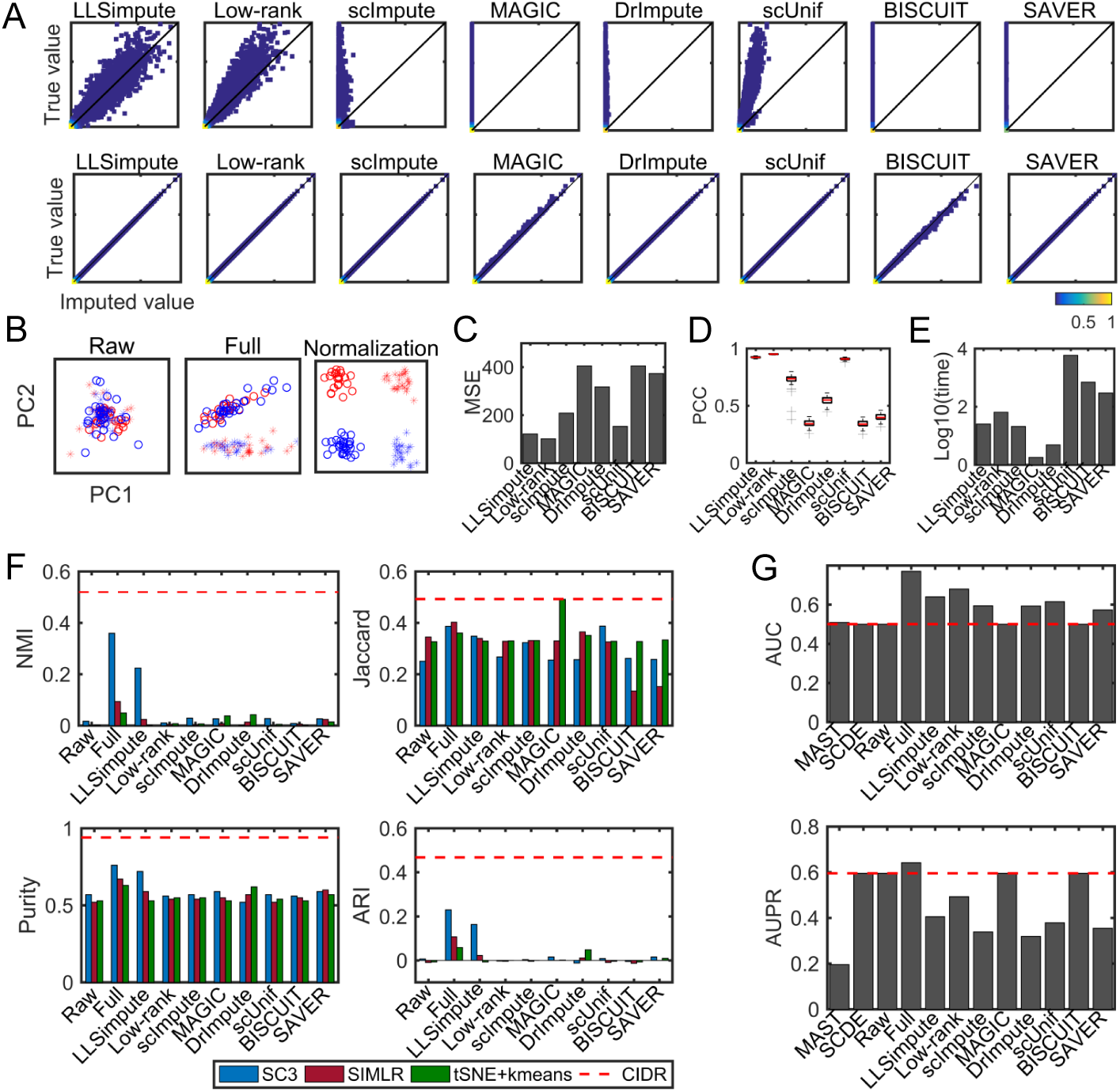
Performance of the eight imputation methods on simulated dataset 5. (A) Density plot of the imputed values versus the original ones in the zero space (top) and the observed non-zero space (bottom), respectively. (B) PCA visualization of the raw data and the original one with groups represented by different colors and batches denoted by different shapes. (C) MSE values of each imputation method. (D) PCC values of all single cell pair computed between the imputed data and the original one. (E) Computational time (seconds) of running each imputation method. (F) Clustering performance of the eight imputation methods on simulated dataset 5. (G) Performance of detecting DEGs.

### 3.3 Imputation methods demonstrate diverse ability in preserving data structures

Proper imputation of dropout values should preserve the underlying data structures. We assessed the imputation methods in several indirect ways including dimension reduction, cell-type clustering, DEG detection and pseudotime trajectory reconstruction. CIDR and ZIFA are two dimension reduction methods, which can address dropout events directly. Compared with CIDR and ZIFA, we visualized the raw data, original data and imputed data by PCA (Figure 5). Low-rank, scImpute, DrImpute, scUnif and SAVER outperform other methods and are consistent with patterns of real data in the first two PC dimensions. Though ZIFA has more divergent clusters than other methods, it is far from real data in the low dimensional space, which might introduce new noise. As the clusters of simulated dataset 3 is not separable in the first two principle components, we also visualized dataset 3 with tSNE. Low-rank, scImpute, DrImpute, scUnif and SAVER still discover different clusters. Interestingly, Low-rank gets the most similar structure with that in the real data in the low-dimensional space (Figure 6B).

**Fig. 5.**
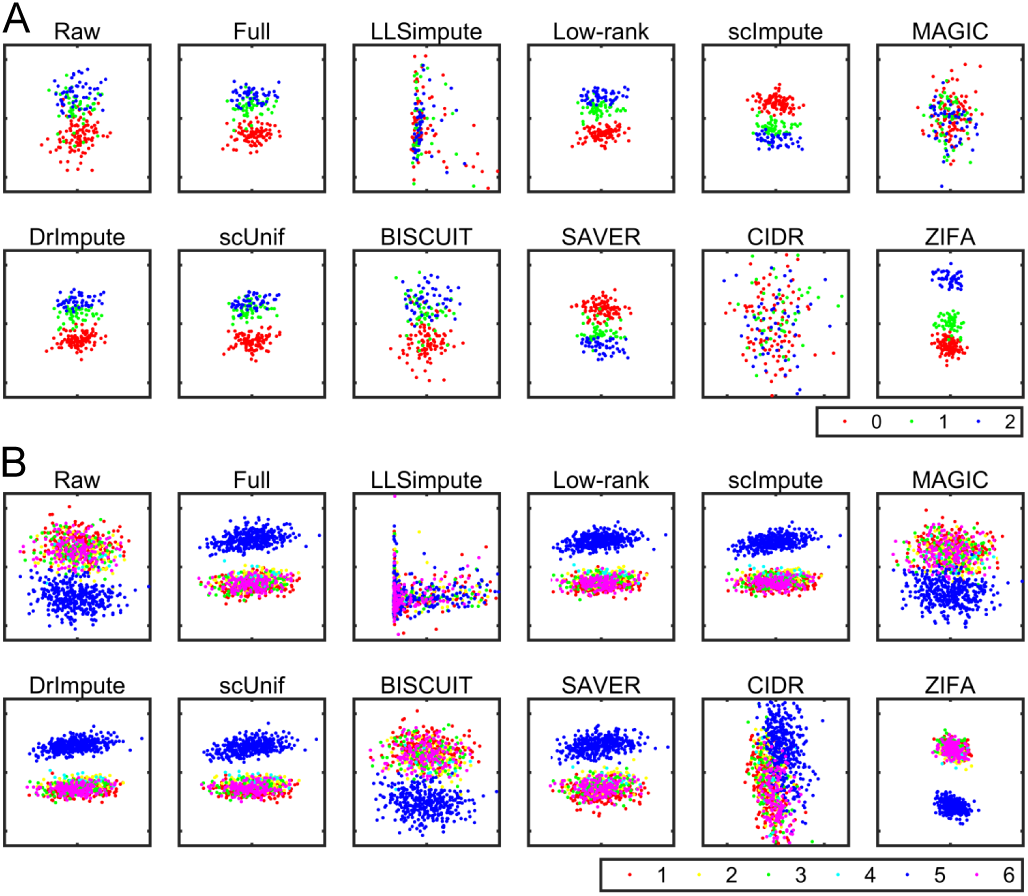
PCA visualization of the reduced dimensions of the eight imputation methods on simulated datasets 2 (A) and 3 (B).

**Fig. 6.**
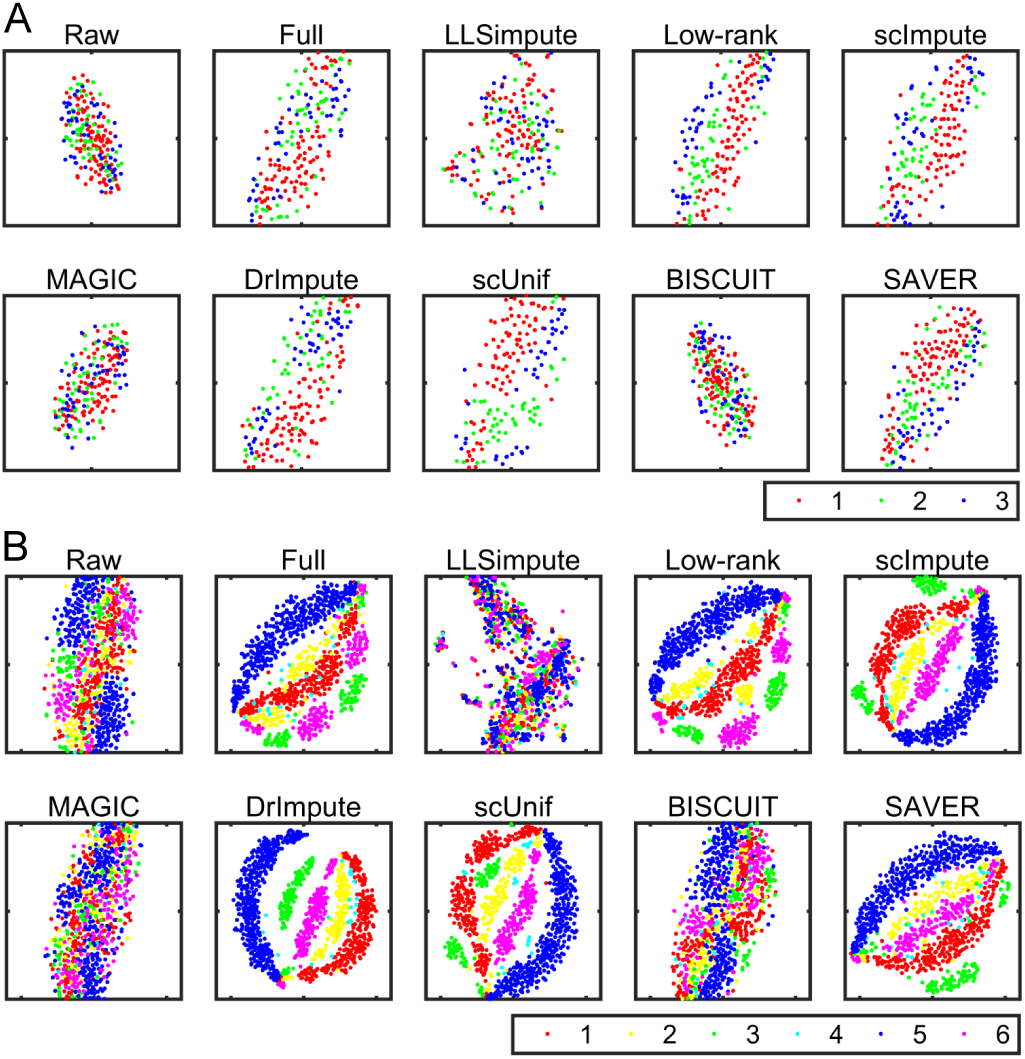
tSNE visualization of the reduced dimensions of the eight imputation methods on simulated datasets 2 (A) and 3 (B).

Identifying cell subpopulations is a key application of scRNA-seq and some clustering methods may fail due to the existence of dropout events. SC3 and SIMLR have been developed for clustering scRNA-seq data, and both of them do not address dropout events directly. We evaluated the effectiveness of these imputation methods with impacts on the clustering performance of SC3, SIMLR and *k*-means with the first two tSNE dimensions (tSNE+kmeans). The clustering performance was assessed by the normalized mutual information (NMI) [32], Jaccard index, purity, and adjusted rand index (ARI). SC3 shows better performance than SIMLR and tSNE+kmeans on simulated datasets 2 and 3. As the first two tSNE components capture little information of the simulated dataset 2 (Figure 6A), tSNE+kmeans has a worse clustering performance. Low-rank, scImpute, DrImpute, scUnif and SAVER improve the clustering performance of SC3 on simulated datasets 2 and 3. In the simulated dataset 2, scUnif and scImpute have the best performance, but scUnif needs cell labels information in advance. Low-rank and DrImpute also have better performance (Figure 7). In the large simulated dataset 3, Low-rank, scImpute, DrImpute, scUnif and SAVER also enhance the clustering performance of SIMLR and tSNE+kmeans. Interestingly, SIMLR and tSNE+kmeans applied to the imputed data by Low-rank, scImpute, DrImpute, scUnif and SAVER have better performance than CIDR (Figure 8). However, CIDR outperforms both of these clustering methods even on the real data on simulated dataset 5 (Figure 4F).

**Fig. 7.**
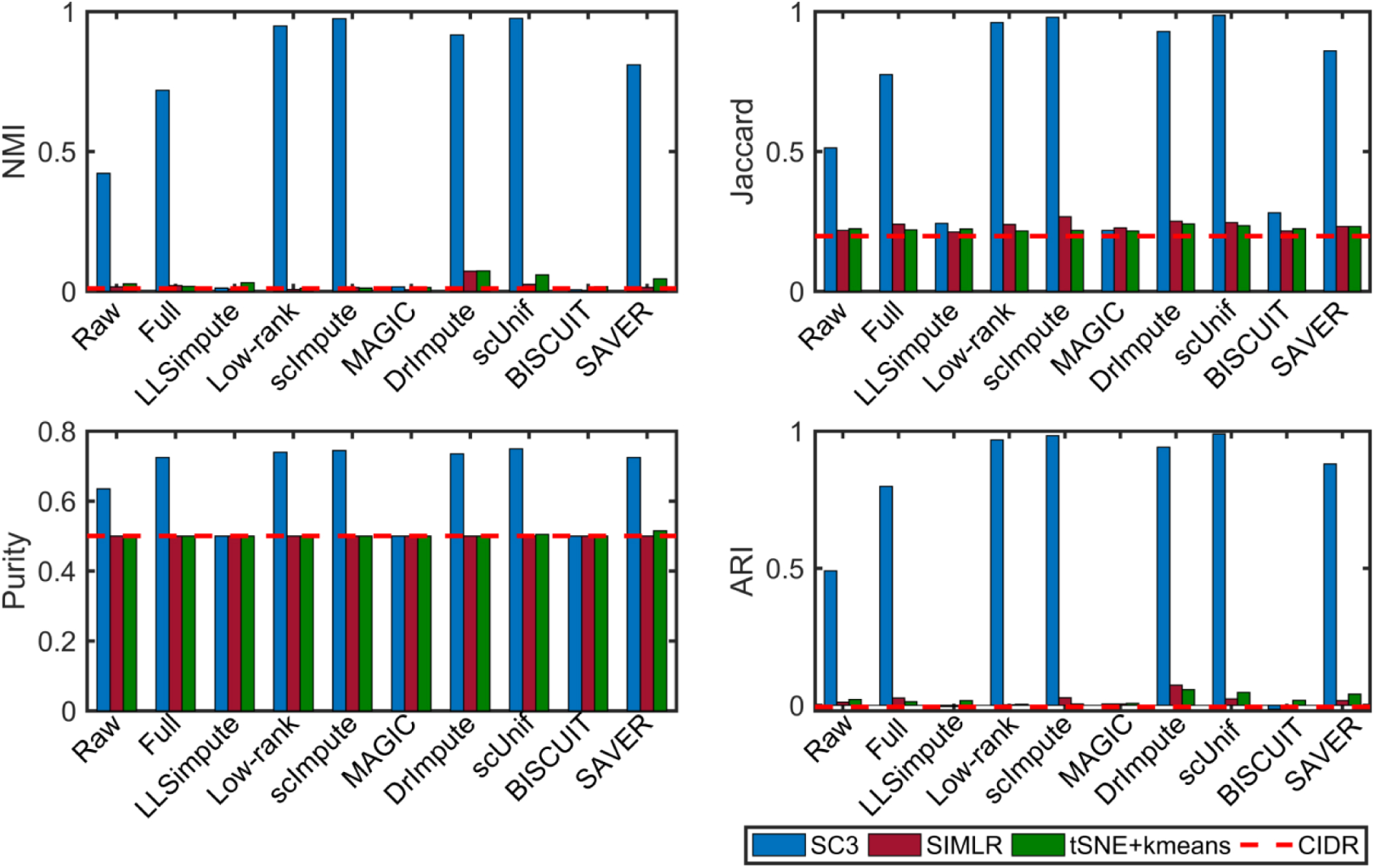
Clustering performance of the eight imputation methods on simulated dataset 2.

**Fig. 8.**
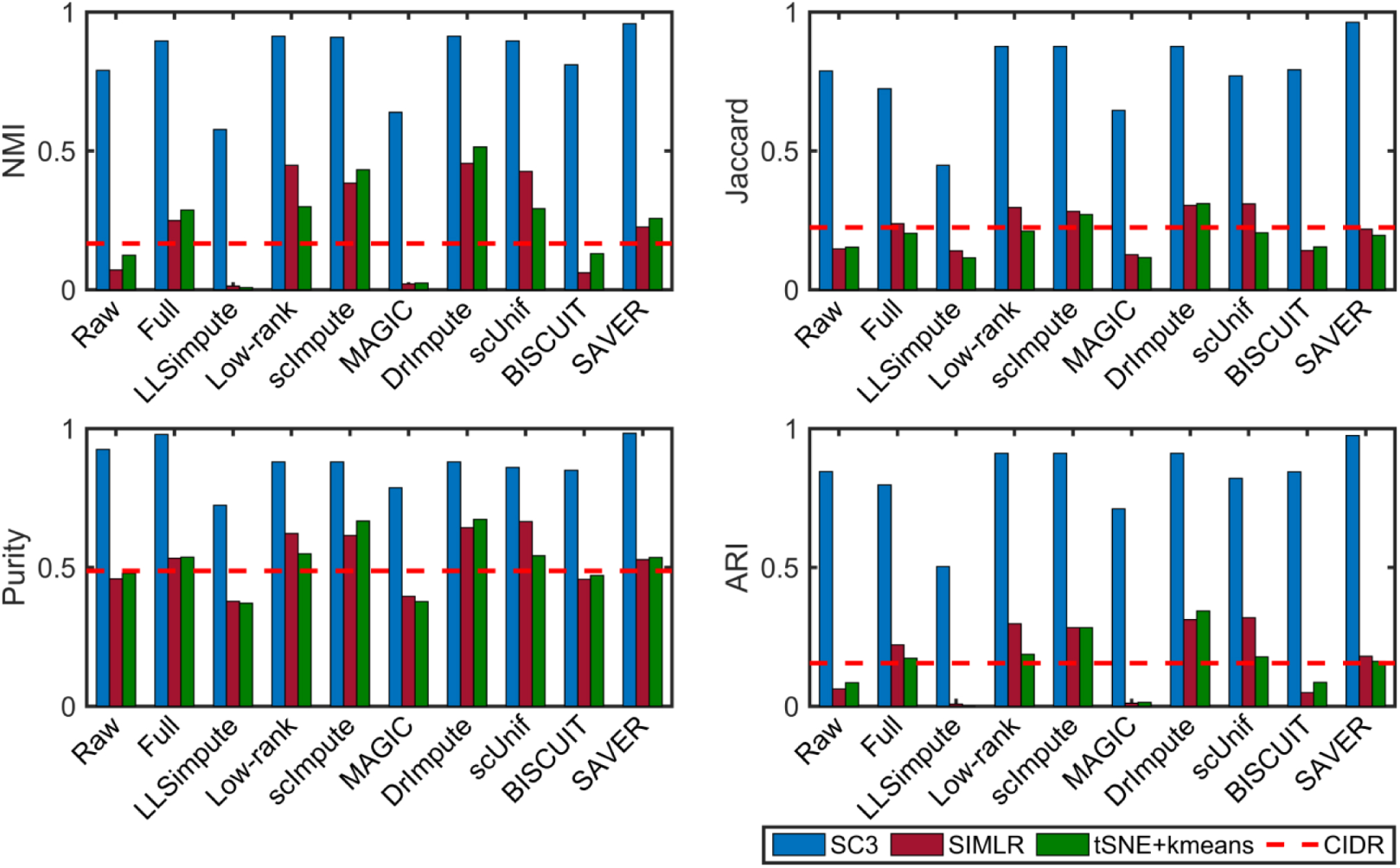
Clustering performance of the eight imputation methods on simulated dataset 3.

To evaluate the robustness of imputation methods, we down-sampled 50 cells at random on simulated dataset 2 with five repetitions. We clustered the down-sampled cells with or without imputing the dropout events by SC3, SIMLR and tSNE+kmeans respectively. The clustering performance show that scUnif has the best robustness, while scImpute is not as good as that of using all cells (Figure 9).

**Fig. 9.**
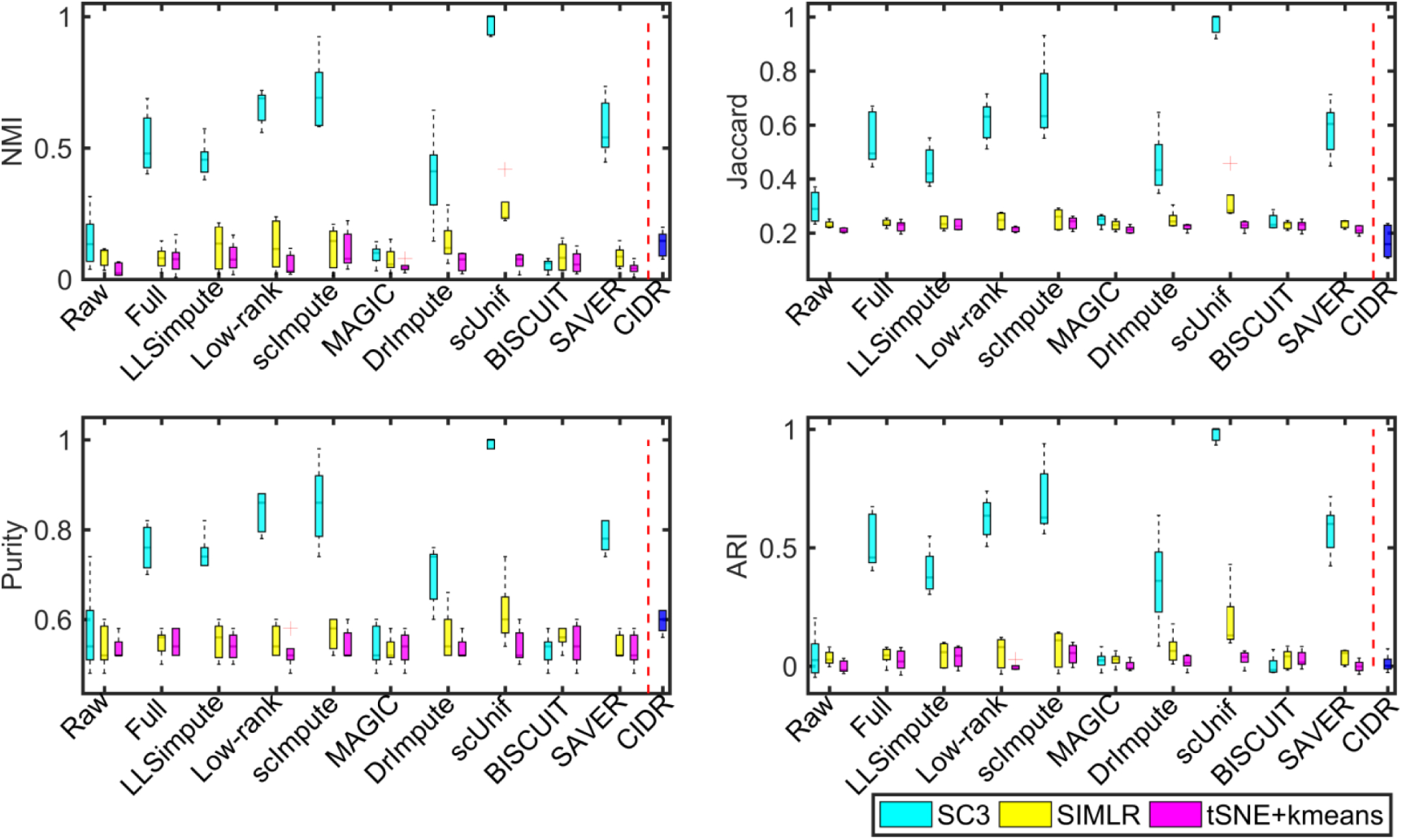
Clustering performance of the eight imputation methods on the down-sampled cells of simulated dataset 2.

Detecting DEGs is also an important downstream analysis of scRNA-seq data. We assessed the recovery power of imputation methods in identifying DEGs from the raw data. We can see that MAST has the worst performance compared to other methods in these two simulated datasets 2 and 3. scImpute, DrImpute and scUnif slightly enhance the performance of edgeR in detecting DEGs, which are better than MAST and SCDE. edgeR has similar performance on raw data with Low-rank, BISCUIT and SAVER on the imputed data. SCDE demonstrate the best AUPR value than other methods (Figure 10). Moreover, the imputation methods have no advantages on improving the performance of edgeR on the simulated dataset 3 (Figure 11). DrImpute and BISCUIT have better performance than other methods, while LLSimpute and MAGIC have the worst performance on simulation dataset 3. Interestingly, imputation methods except MAGIC and BISCUIT enhance the sensitivity of edgeR in detecting DEGs, while these methods have smaller AUPR values than those of MAGIC and BISCUIT (Figure 4G).

**Fig. 10.**
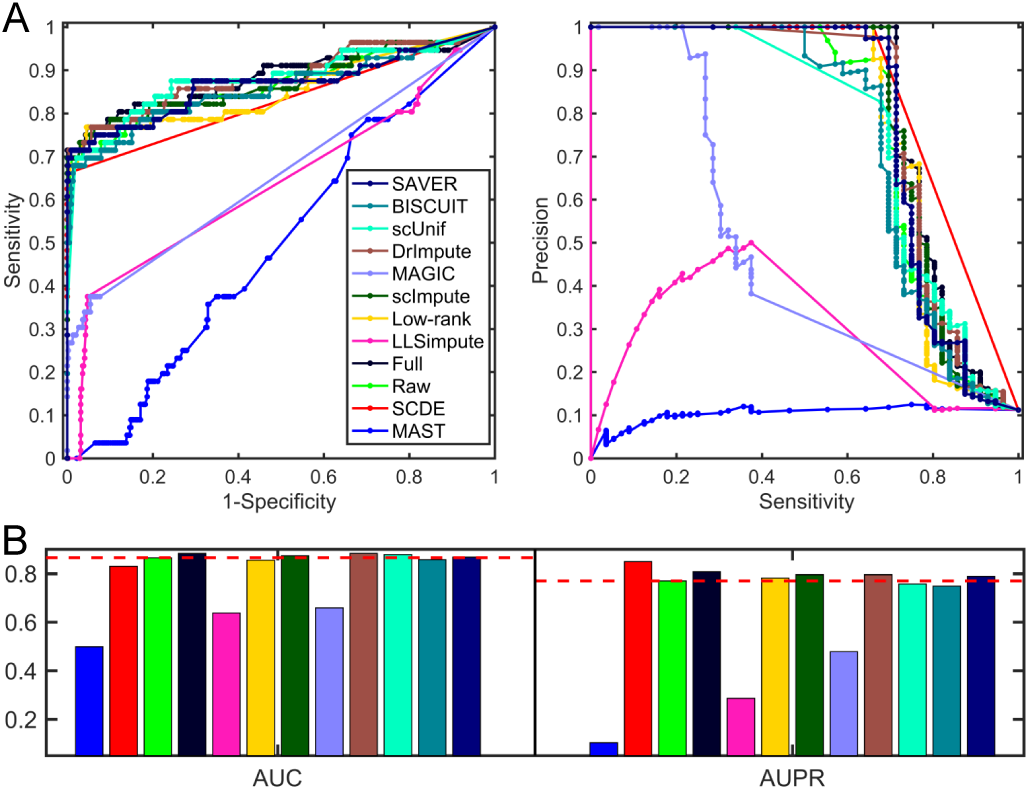
Performance of detecting DEGs on simulated dataset 2.

**Fig. 11.**
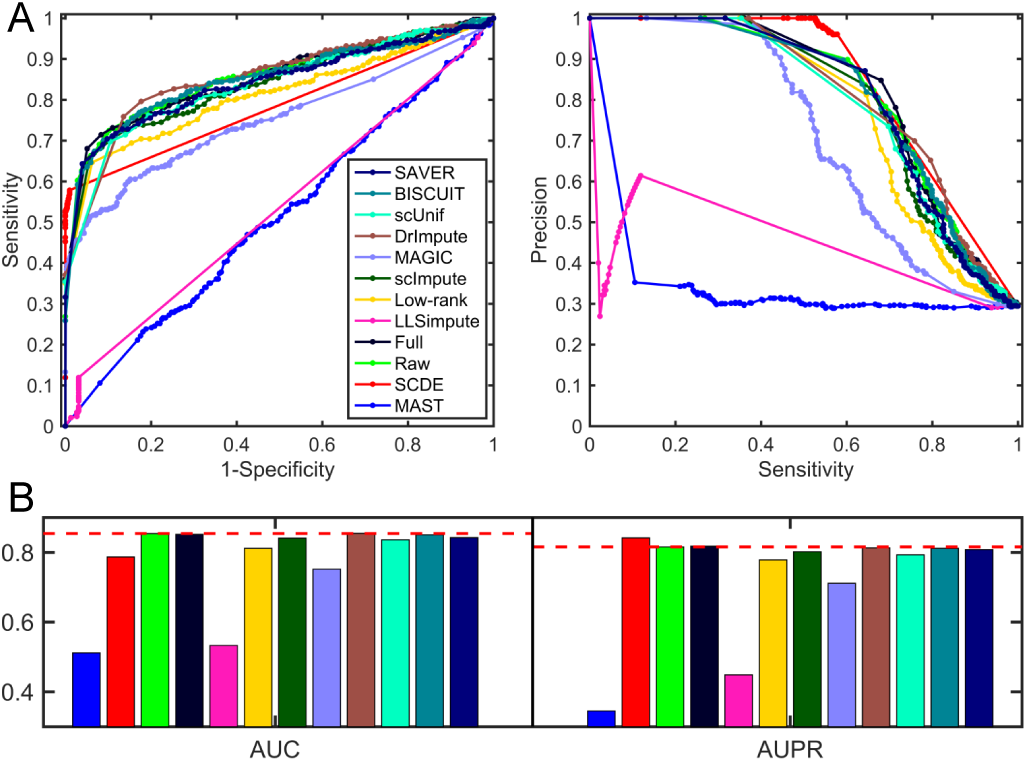
Performance of detecting DEGs on simulated dataset 3.

scRNA sequencing has already shown its power in reconstructing developmental trajectories [33]. We employed the simulated dataset 4 with a path and no branches to compare the impacts of imputation methods on inferring pseudotime order. As LLSimput imputed data does not satisfy the requirement of Monocle 2, we did not show the pseudotemporal order of it. scImpute and SAVER have more consistent trajectories with that constructed in the original data (Figure 12). The measurable indicator of order conformity (named as order correlation) is defined as *C*/(*N_c_* + *C*), where *C* represents the number of similar orders between the pseudotime and gold standard orders, and *N_c_* denotes the number of dissimilar orders. scImpute, DrImpute and SAVER improve the power of Monocle 2 in ordering cells along a trajectory in terms of this index. We down-sampled 50 cells randomly for five repetitions, imputed them by these imputation methods and inferred trajectories with or without imputing the dropout events. We can see that Low-rank has the largest order correlations than those of other methods, enhancing the power of Monocle 2 by imputing the dropout events. Moreover, MAGIC has the worse performance when we randomly selected some cells to infer a trajectory.

**Fig. 12.**
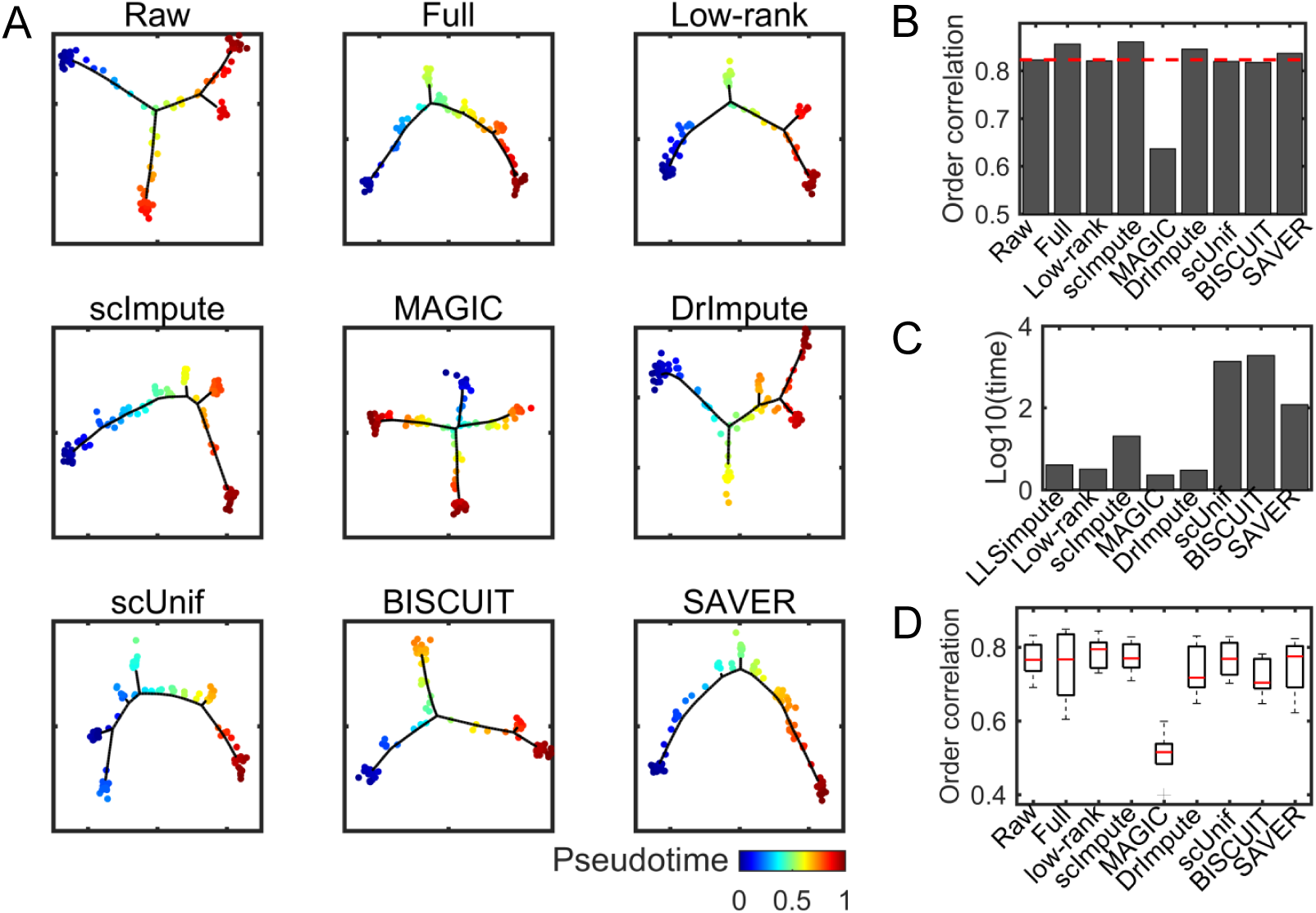
Performance of reconstructing pseudotime order of scRNA-seq data using imputation methods or not on simulated dataset 4. (A) Visualization of the inferred trajectory using each method. Each dot represents a cell. Cells with higher values are in more differentiated states. (B) Order correlation of pseudotime inferred from Monocle 2 on the raw data, original data, and imputed data with golden standard pseudotime. (C) Computational time (seconds) of running each imputation method. (D) Boxplot of order correlation of five repetitions with 50 cells.

In summary, Low-rank, scImpute, DrImpute, scUnif and SAVER have better performance in dimension reduction and cell clustering, which are even better than CIDR. scImpute, DrImpute and SAVER improves the performance of Monocle 2 in reconstructing pseudotime. However, these imputation methods have no significant improvement for edgeR in detecting DEGs. Only scImpute, DrImpute and scUnif slightly enhance this performance on simulated dataset 2.

### 3.4 Imputation methods provide more potential DEGs

In the mECS real data, the 182 mESC cells consist of 59 cells in G1 phase, 58 cells in S phase and 65 cells in G2/M phase. Firstly, BISCUIT and SAVER impute zeros with near zero values, while LLSimpute and scImpute impute zeros with relatively large values. We compared true values with recovered ones by Low-rank, MAGIC, BISCUIT and SAVER, which will change the values in the observed space in principle (Figure 13B). Low-rank and SAVER recover the observed values well, while MAGIC and BISCUIT change the observed ones in some degree. However, MAGIC improves the clustering performance of SC3. BISCUIT enhances the power of SC3 in clustering slightly. tSNE+kmeans has better clustering performance on DrImpute imputed data than on the raw data, and even better than CIDR, which addresses dropout events directly. edgeR on LLSimpute imputed data tends to treat each genes as a DEG, which is fallacious (Figure 13D). There are 332 DEGs by SCDE with *q*-value < 0.05, which are included in the DEG set of edgeR on Low-rank imputed data. We downloaded mouse cell cycle stage-specific maker genes from a previous study [31], which includes 43 (31) and 54 (51) marker genes of G1/S and G2/M (in the processed data) respectively.

**Fig. 13.**
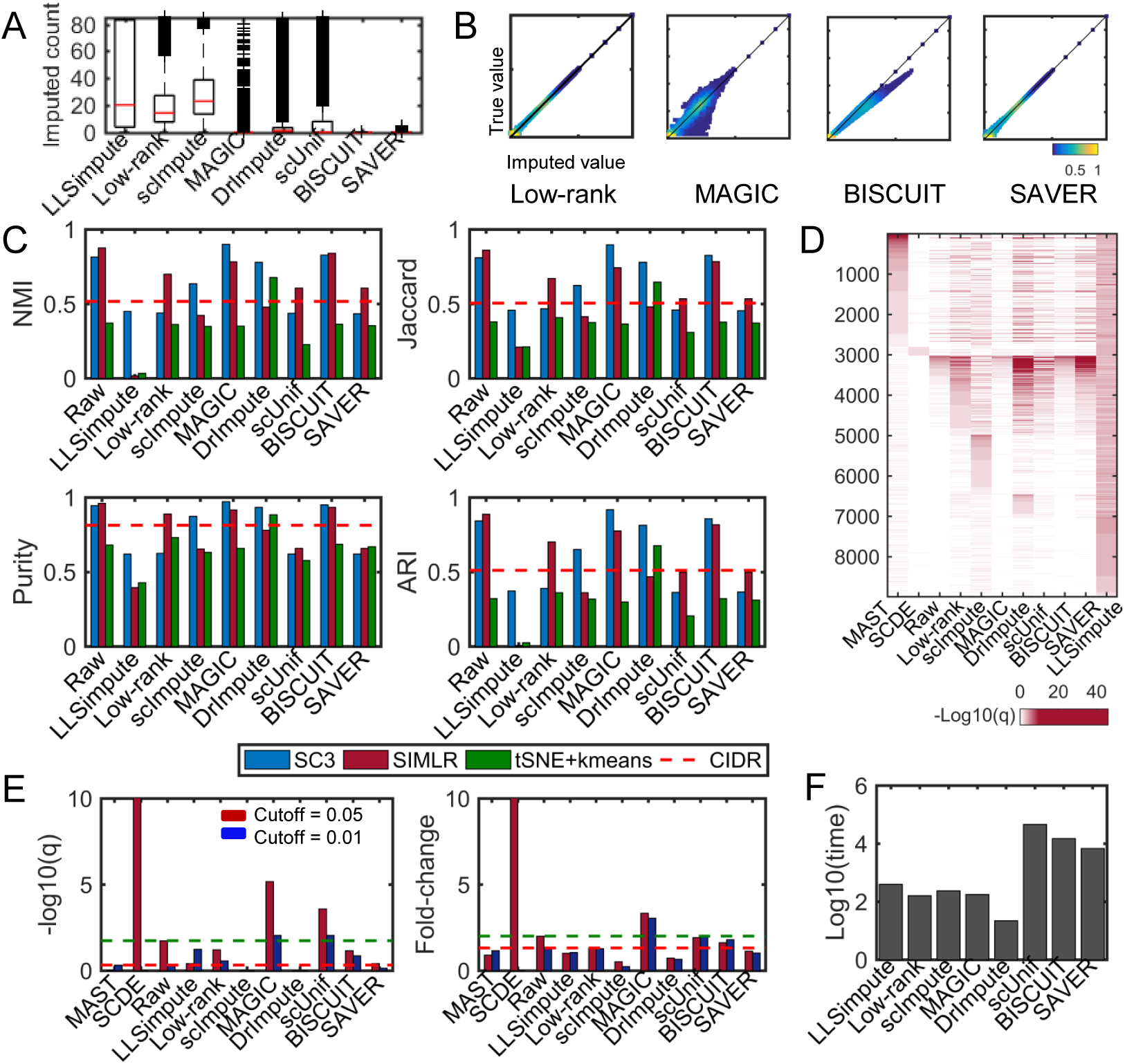
Performance of the eight imputation methods on the mESC data. (A) Boxplot of the imputed values of each method on the zero space. (B) Density plot of the recovered values versus original ones in the observed non-zero space using Low-rank, MAGIC, BISCUIT and SAVER with *y*-axis representing log-transformed observed non-zero values and *x*-axis denoting log-transformed recovered values. (C) Clustering performance of CIDR on raw data and SC3, SIMLR, tSNE+*k*-means on raw data and imputed data in terms of NMI, Jaccard, Purity and ARI respectively. (D) Heatmap of −log10(*q*) value of MAST, SCDE on raw data and edgeR on raw and imputed data for detecting DEGs. (E) Barplot of −log10(q) and fold-change of Fishers exact test on the enrichment analysis of 82 G1/S, G2/M maker genes. (F) Computational time (seconds) of running each imputation method.

The DEGs of SCDE is significantly enriched with the G1/S and G2/M marker genes using Fisher’s exact test with FDR < 0.05. However, only *PLK1* is regarded as DEG by SCDE with FDR < 0.01. The activity of *PLK1* is indeed regulated by cell cycle, which is in low activity during interphase but high during mitosis [34]. MAGIC enhances the enrichment of marker genes with higher fold-changes (Figure 13E). There are six G2/M marker genes detected by eageR on MAGIC imputed data but ignored by SCDE. These genes indeed have higher count level in G2 cells recovered by MAGIC (Figure 14). The Bayesian-based imputation methods are still more time-consuming than other methods (Figure 13F).

**Fig. 14.**
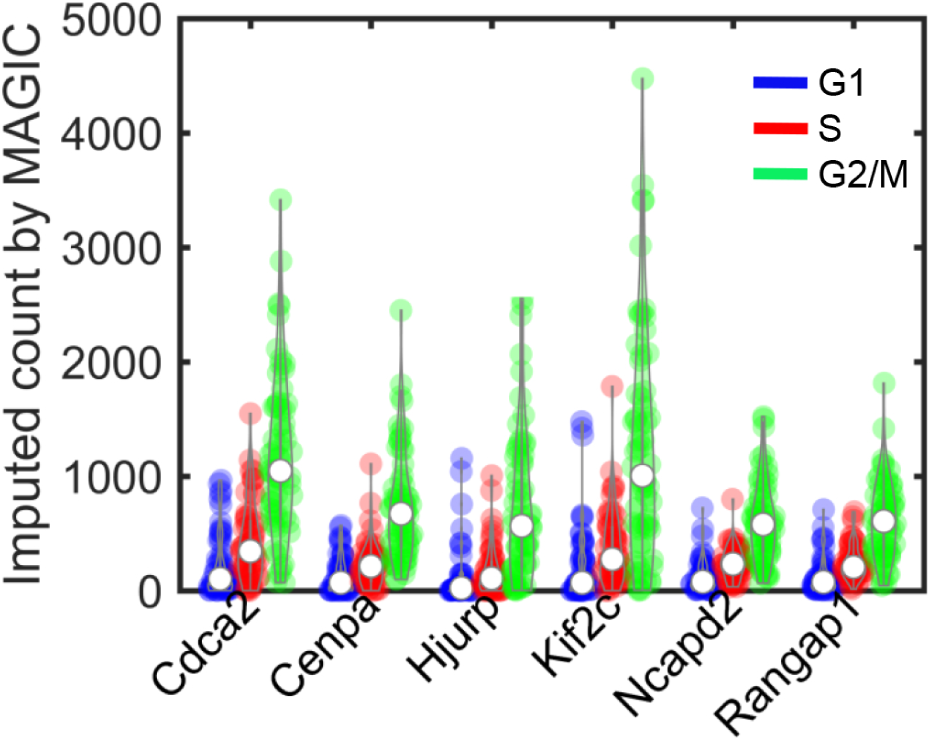
Violinplot of the imputed count values of G2/M marker genes, which are detected by edgeR using MAGIC imputed data but ignored by SCDE.

### 3.5 Imputation methods improve the reconstruction of epidermal differentiation process

The great regenerative capacity of murine epidermis and its appendages enable it to be an invaluable model system for stem cell biology. Recently, the cellular heterogeneity of the adult mouse epidermis has been examined using scRNA-seq. The pseudotemporal order of IFE cells has been obtained by a minimum spanning tree-based method in tSNE space [31]. Applied imputation methods to this dataset, we can see that BISCUIT still imputes zeros with near zero values in this data. MAGIC and SAVER recover non-zero values with large deviation, while BISCUIT recover non-zero values well, indicating that there is small technical variation detected by BISCUIT in this data (Figure 15A and 15B).

**Fig. 15.**
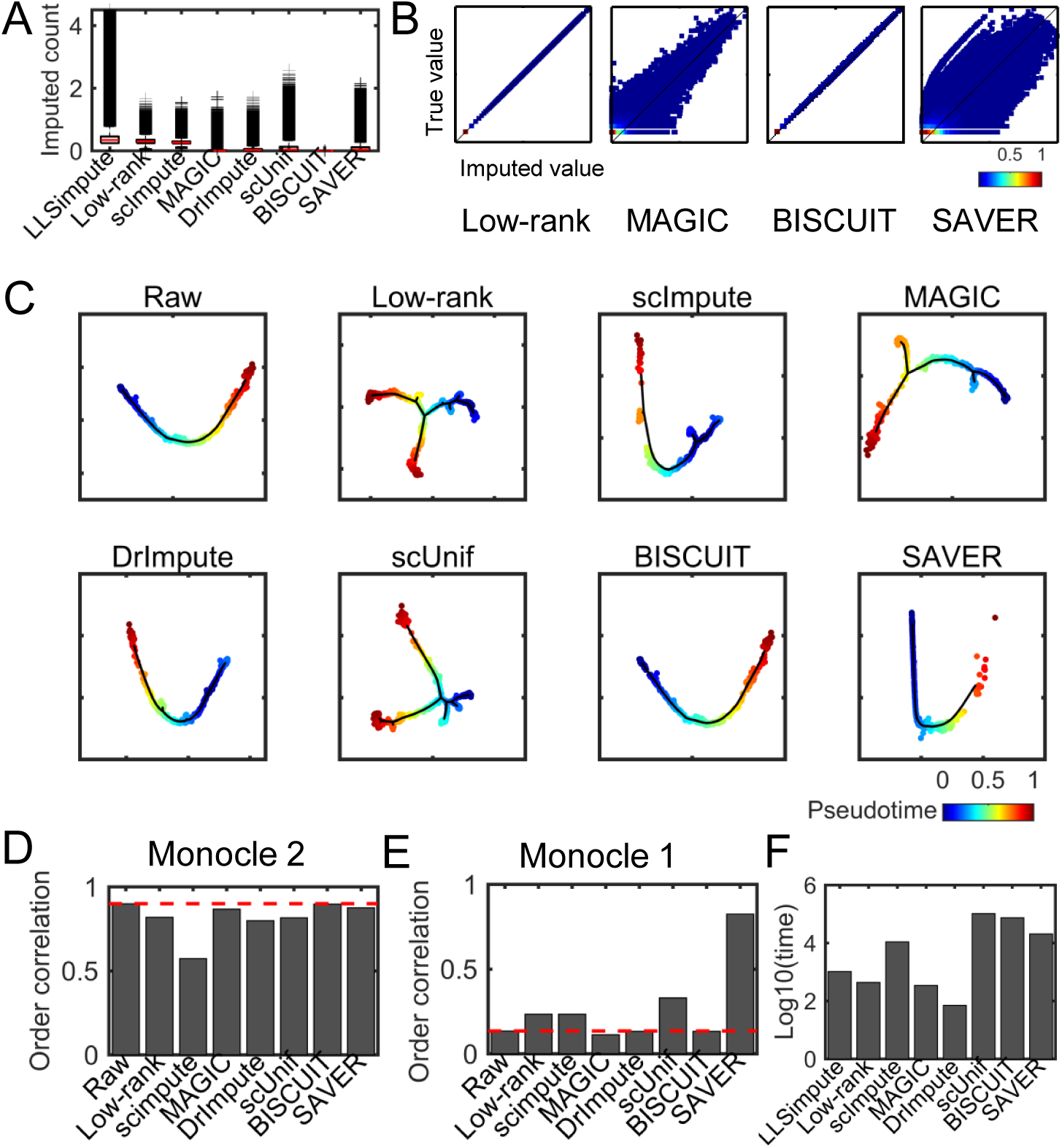
Pseudotime reconstruction of IFE data. (A) Boxplot of the log-transformed imputed values of the eight imputation methods on the zero space. (B) Density plot of the recovered values versus the original ones in the observed non-zero space using Low-rank, MAGIC, BISCUIT and SAVER with *y*-axis representing log-transformed true values and *x*-axis denoting log-transformed recovered values. (C) Visualization of the inferred trajectory of each method. Each dot represents a cell. Cells with higher values are in more differentiated states. (D) Order correlation of pseudotime inferred from Monocle 2 on the raw data and imputed data with golden standard pseudotime. (E) Order correlation of pseudotime inferred from Monocle 1 on the raw data and imputed data with golden standard pseudotime. (F) Computational time (seconds) of running each imputation method.

We inferred the pseudotime of individual cells by Monocle 1 [23] and Monocle 2 [24] on the raw data and imputed data respectively. The transcribed repetitive elements (i.e., gene name stated with “*r*_”) were removed from the gene list. The trajectory reconstructed on the raw data and imputed data by these imputation methods except Low-rank and scUnif are more consistent with the one inferred by the spanning tree model visually [31]. Based on the order correlation, MAGIC, BISCUIT and SAVER preserve the differentiation direction well, which starts from IFE basal cells to IFE differentiated cells, then arrives at IFE keratinized layer (Figure 15D). However, any imputation methods cannot enhance the performance of Monocle 2 due to its efficiency. Interestingly, Monocle 1 is not as effective as Monocle 2 on the raw data. SAVER, Low-rank, scUnif and scImpute improve the ability of Monocle 1 clearly (Figure 15E). Therefore, the impact of imputing dropout events may rely on the downstream analysis method. MAGIC and BISCUIT have more reliable IFE differentiation process as the pseudotime order of the known basal maker (Krt14), mature marker (Krt10), terminally differentiated cell stage marker (Lor) and a transient marker (Mt4) gradually vary along the trajectory within each cluster than those of other imputation methods (Figure 16).

**Fig. 16.**
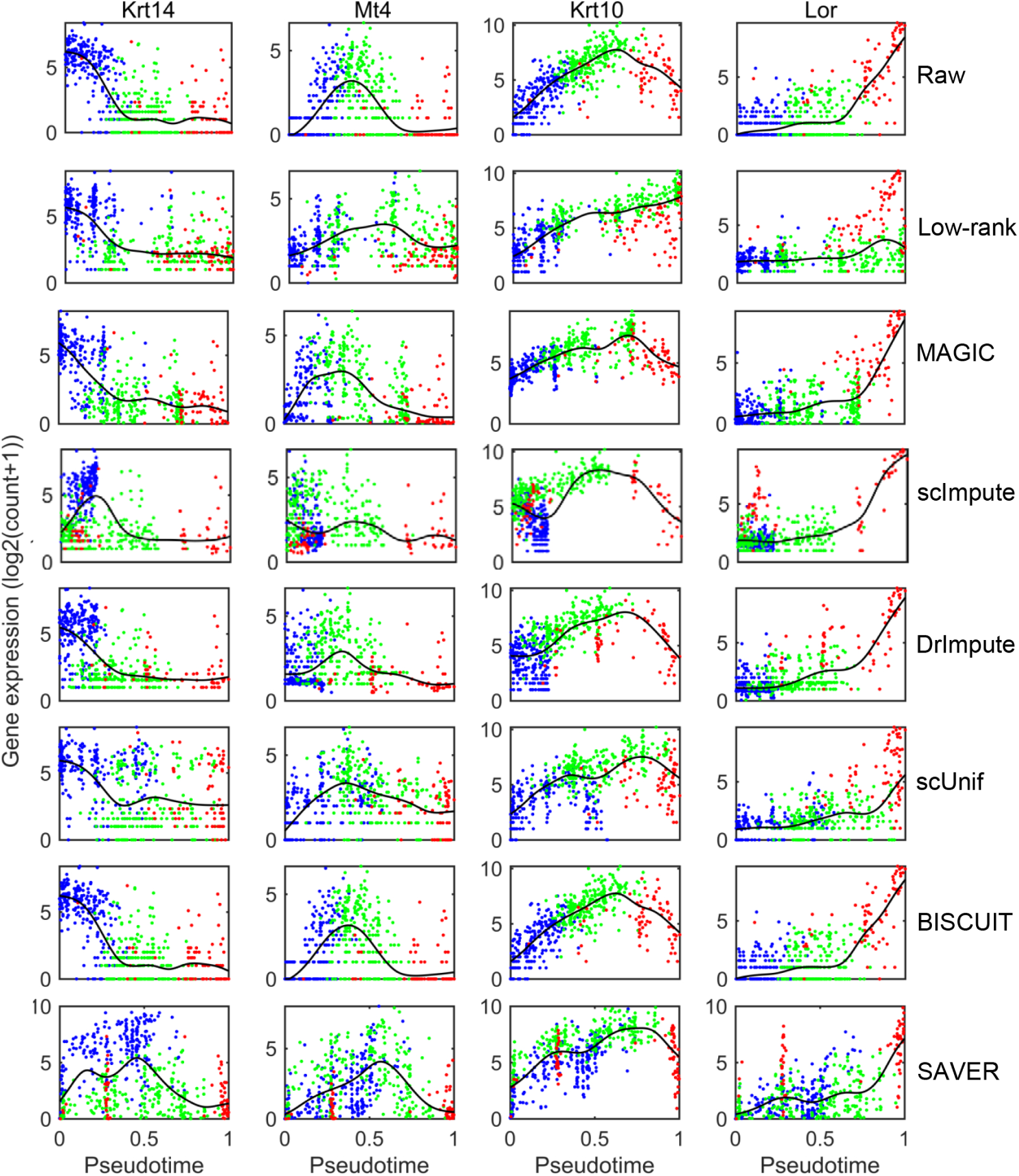
Validation of pseudotemporal ordering of IFE cells with four marker genes Krt14, Krt10, Lor and Mt4 using the raw data and imputed data by Monocle respectively. Each dot represents a cell. The IFE basal cells, IFE differentiated cells and IFE keratinized layer are denoted by blue, green and red colors respectively. The gene expression of these markers were fitted with cubic smoothing splines.

## 4 Conclusion and discussion

The main goal of this study is to provide a straightforward and thorough comparison on the imputation methods for scRNA-seq data. We systematically evaluated eight imputation methods including two for general incomplete data and six specially designed for scRNA-seq data from multiple angles. We summarized the impacts of eight imputation methods on the simulated and real datasets (Table 3). Firstly, LLSimpute designed for the bulk-RNAseq data performs well in a homogenous cell population, but it fails when the data shows large heterogeneity and sparsity, which are two key characteristics of scRNA-seq data. Low-rank also performs well in datasets 1 and 5. It is not affected by batch effect applying to all genes. Secondly, scImpute and DrImpute recover the data well in simulated datasets. However, they fail on the data with less collinearity (e.g., mESC data). Thirdly, simulation study illustrates that BISCUIT and SAVER tend to impute the dropout events with near zero values. MAGIC and BISCUIT recover non-zero values with large fluctuations. MAGIC shows better performance to help to detect biomarkers in mouse mESC data. MAGIC designed based on a Markov affinity-based graph could capture the gradual variation of genes. Therefore, it enhances the performance of Monocle 2 to reconstruct marker genes expression change along differentiation process. However, it fails to improve the ability of Monocle 1. These results demonstrate that the impacts of imputing dropout events on downstream analysis depend on the analysis methods.

**TABLE 3.**
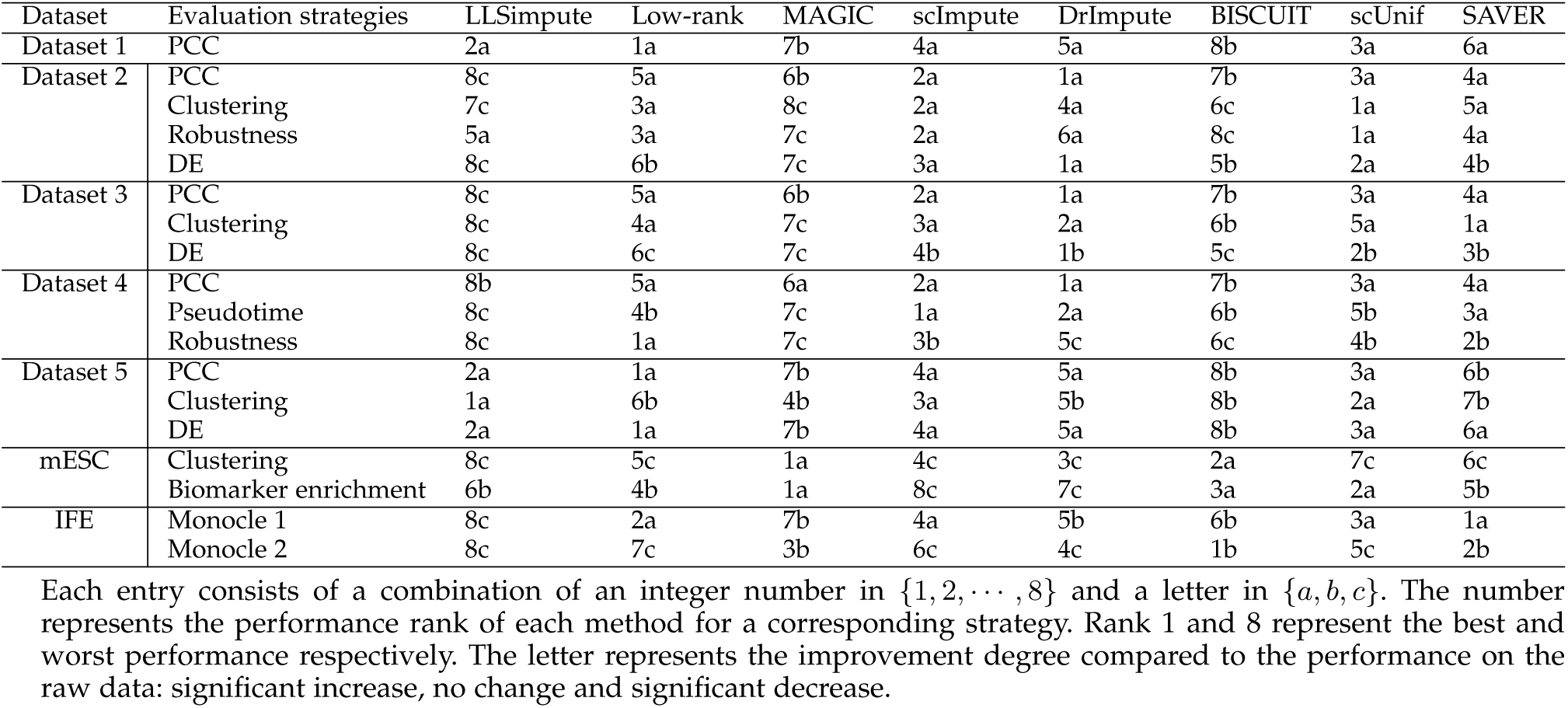
Summary of systematic evaluation of the eight methods using both simulated and real datasets

Extensive studies highlight that there is no one method performs the best in all situations. Current methods still have some defects such as scalability, robustness and applicability in some situations. With the rapid generation of large-scale scRNA-seq data, imputation of dropout events is becoming a basic and routine step in scRNA-seq data analysis. Therefore, efficient methods and powerful tools for imputation are urgently needed at present. Moreover, efficient information from genes such as co-expressed networks should be used in future studies.

## Acknowledgment

Shihua Zhang is the corresponding author of this paper. This work has been supported by the National Natural Science Foundation of China [No. 61422309, 61379092, 61621003 and 11661141019]; the Strategic Priority Research Program of the Chinese Academy of Sciences (CAS) [No. XDB13040600], the Key Research Program of the Chinese Academy of Sciences [No. KFZD-SW-219] and CAS Frontier Science Research Key Project for Top Young Scientist [No. QYZDB-SSW-SYS008].

